# Kinetic mechanism for fidelity of CRISPR-Cas9 variants

**DOI:** 10.1101/2025.01.27.634389

**Authors:** Andrew D. Hecht, Oleg A. Igoshin

## Abstract

CRISPR-Cas9 is a nuclease creating DNA breaks at sites with sufficient complementarity to the RNA guide. Notably, Cas9 does not require exact RNA-DNA complementarity and can cleave off-target sequences. Various high-accuracy Cas9 variants have been developed, but the precise mechanism of how these variants achieve higher accuracy remains unclear. Here, we develop a kinetic model of Cas9 substrate selection and cleavage. We parameterize the model using datasets available in the literature, including both high-throughput substrate binding and cleavage data and Förster resonance energy transfer measurements of the Cas9 HNH domain transitions. Based on the observed transition statistics, we predict that the Cas9 substrate recognition and cleavage mechanism must allow for HNH domain transitions independent of substrate binding. Additionally, we show that the enhancement in Cas9 substrate specificity must be due to changes in kinetics rather than changes in substrate binding affinities. Furthermore, the fitted model produces quantitatively realistic cleavage error predictions for substrates with protospacer adjacent motif (PAM)-distal mismatches. Finally, we use our model to identify kinetic parameters for HNH domain transitions that can be perturbed to enable high-accuracy cleavage while maintaining cleavage speeds. Our results refine the biophysical mechanism of Cas9 cleavage to inform future routes for its engineering.

## I. INTRODUCTION

CRISPR-Cas9 is an RNA-guided DNA nuclease derived from bacteria [13]. Valid DNA target sites for Cas9 require the presence of a short DNA sequence known as a protospacer adjacent motif (PAM) that is recognized via protein-DNA interactions [12]. After initial PAM binding, the adjacent DNA is unwound to allow the Cas9 guide RNA to interrogate the local sequence, causing the formation of an RNA-DNA hybrid known as an R-loop. The Cas9 RuvC and HNH nuclease domains then become catalytically active following full R-loop formation and can cleave the backbone of both DNA strands.

Cas9 is capable of cleaving at target sites even in the presence of DNA-RNA mismatches [5, 11]. Generally, mismatches at the PAM-distal end of the target are better tolerated than PAM-proximal (so-called “seed region”) mismatches. Inappropriate cleavage at such off-target sites can cause DNA base insertion/deletion (indel) mutations as well as genomic-scale damage in the form of large deletions [2], which both present significant safety concerns for clinical applications of Cas9-based technologies. Accordingly, multiple efforts have been made to develop high-fidelity CRISPR-Cas9 platforms [1, 6, 17, 28, 29].

The high-fidelity variants developed by Slaymaker and Kleinstiver were built under the “excess energy” hypothesis [17, 28]. Under this hypothesis, the Cas9-DNA binding interaction is thought to be more energetically favorable than necessary to achieve catalytic activation and subsequent DNA cleavage. Thus, the bound complex would be destabilized by removing Cas9 residues that interact non-specifically with the DNA backbone via electrostatics. This destabilization then increases the probability that RNA-DNA mismatches would disrupt enzymatic activation. The resulting variants were indeed shown to achieve higher substrate selection fidelity than the wild-type Cas9 variant [17, 28]. However, there was no significant difference in substrate binding affinity compared to WT-Cas9 when bound to either fully matched or partially mismatched sequences [1]. Therefore, an alternative mechanism could be required to explain the improved specificity.

In an effort to explain the specificity enhancement observed in the high-fidelity variants, single-molecule Förster resonance energy transfer (smFRET) experiments examined the effects of the dynamics of the Cas9 HNH domain on substrate selection specificity [1, 3]. It was found that the Cas9 HNH domain can adopt distinct conformational states, including an intermediate FRET “checkpoint” conformation that appears to play a role in rejected distally-mismatched substrates. The HNH domain dynamics of the high-fidelity Cas9 variants were found to be perturbed relative to the wild-type variant, with the high-fidelity variants being significantly more sensitive to the presence of PAM-distal mismatches. However, it remains unclear whether the dynamics of the HNH domain directly control cleavage specificity or if the changes in the dynamics are a result of some other process that underlies the change in specificity (e.g. R-loop formation and collapse).

In order to develop our understanding of Cas9 substrate selection, we developed a biophysical model of target site selection and cleavage. Our model proposes a kinetic scheme to model Cas9 transitions with parameters estimated based on fitting of multiple datasets in the literature, including single-molecule FRET measurements of the Cas9 HNH domain [1, 3] and high-throughput measurements of Cas9-DNA binding and cleavage [14]. We use the parameterized model to examine the relative importance of key Cas9 target selection processes, including R-loop formation and HNH domain movements, toward off-target rejection. Finally, we use our model to examine trade-offs between on-target cleavage speed and off-target specificity and show that there are few intrinsic trade-offs preventing the development of high-speed, high-specificity Cas9 variants.

## II. RESULTS

### A. Changes in binding are insufficient to explain high-fidelity Cas9 accuracy

One of the simplest ways to model CRISPR-Cas9 substrate selection is as a linear (1D) process (Fig. 1A) with two possible branches corresponding to cleaving either a correct, on-target substrate (R) or an incorrect, off-target substrate (W). Such a model is a special case of the Shvets-Kolomeisky model of Cas9 [25] where there is only a single off-target site available. Binding to either substrate is a two-step process, starting with initial binding to the PAM site, which is then followed by the formation of a full R-loop. Once the R-loop is fully formed, Cas9 can cleave the bound substrate to form a cleaved DNA product. The substrate selection error *η* can then be defined as a ratio of probabilities *η* = *π*_*W*_/*π*_*R*_, where *π*_*R/W*_ are the conditional probabilities of cleaving one substrate before the other, given that the system starts in state *S*_0,1_ at the initial time *t* = 0.

**Figure 1.**
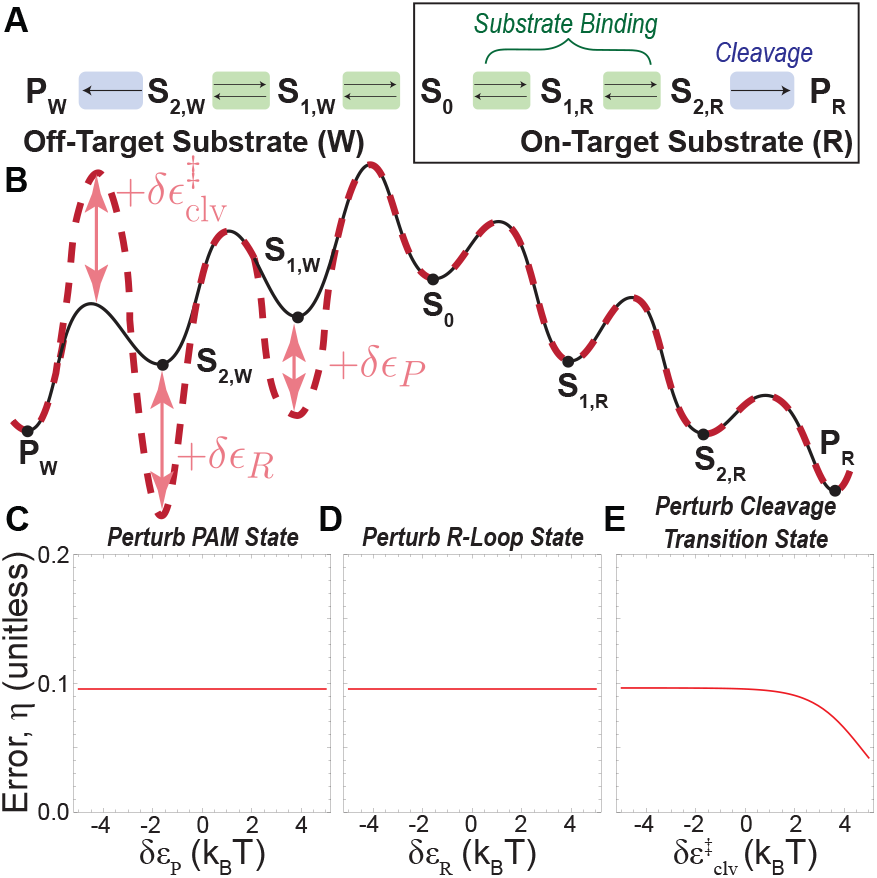
Cas9 selection between an on-target and off-target substrate can be represented by a simple model with two bound states for each substrate (A). The bound states correspond to an initial PAM-bound state and a fully bound, R-loop state. Amino acid substitutions, such as those involved in developing high-fidelity Cas9 variants, can affect Cas9 function by modulating the underlying free-energy landscape (B). The WT-Cas9 free-energy landscape (black) could be perturbed to yield the high-fidelity landscape (red) by changing either the stabilities of individual states (i.e., binding energies) or the barriers between states, which correspond to activation energies. Computing the cleavage error for such a model we see that changes in the stability of states, such as destabilizing the off-target PAM state (C) or the R-loop state (D) are insufficient to affect Cas9 specificity. Only changes in transition state activation energies, such as that for cleaving the off-target substrate (E), can affect specificity.

Thermodynamically and kinetically, this model type can be represented by a free energy landscape (Fig. 1B) in which the individual states of the model correspond to potential wells. Adjacent states are separated by free-energy barriers, with maxima representing transition state energies for transitions between states. In this landscape, two types of perturbations are possible: changes in the energies of the stable intermediate states (*ϵ*_*i*_) or changes in the transition state energies 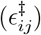.

Previous theoretical work has shown that only perturbations in transition state energies can affect the error of biological substrate selection processes; in other words, the error is under kinetic control [19]. The result holds for the model of Cas9 in Fig. 1A: our calculations demonstrate that decreasing the energy of the off-target R-loop or PAM-bound states relative to the on-target substrate (i.e., as increasing their relative stability) does not change the error (Fig. 1C,D). In contrast, if the substrate cleavage transition-state energy for the off-target substrate is increased, the error decreases substantially (Fig. 1E). Therefore, we conclude that even if the high-fidelity mutations affected the Cas9-DNA binding affinity, it would not cause any change in substrate selection accuracy.

### B. HNH domain transition statistics imply that the energy landscape is multi-dimensional

Experimental single-molecule FRET studies of Cas9 identified three distinct configurational states corresponding to low, mid, and high FRET conformations [1, 3]. Each conformation represents the Cas9 HNH domain moving from the inactive (low-FRET) state towards the catalytically active (high-FRET) state (Fig. 2A).

**Figure 2.**
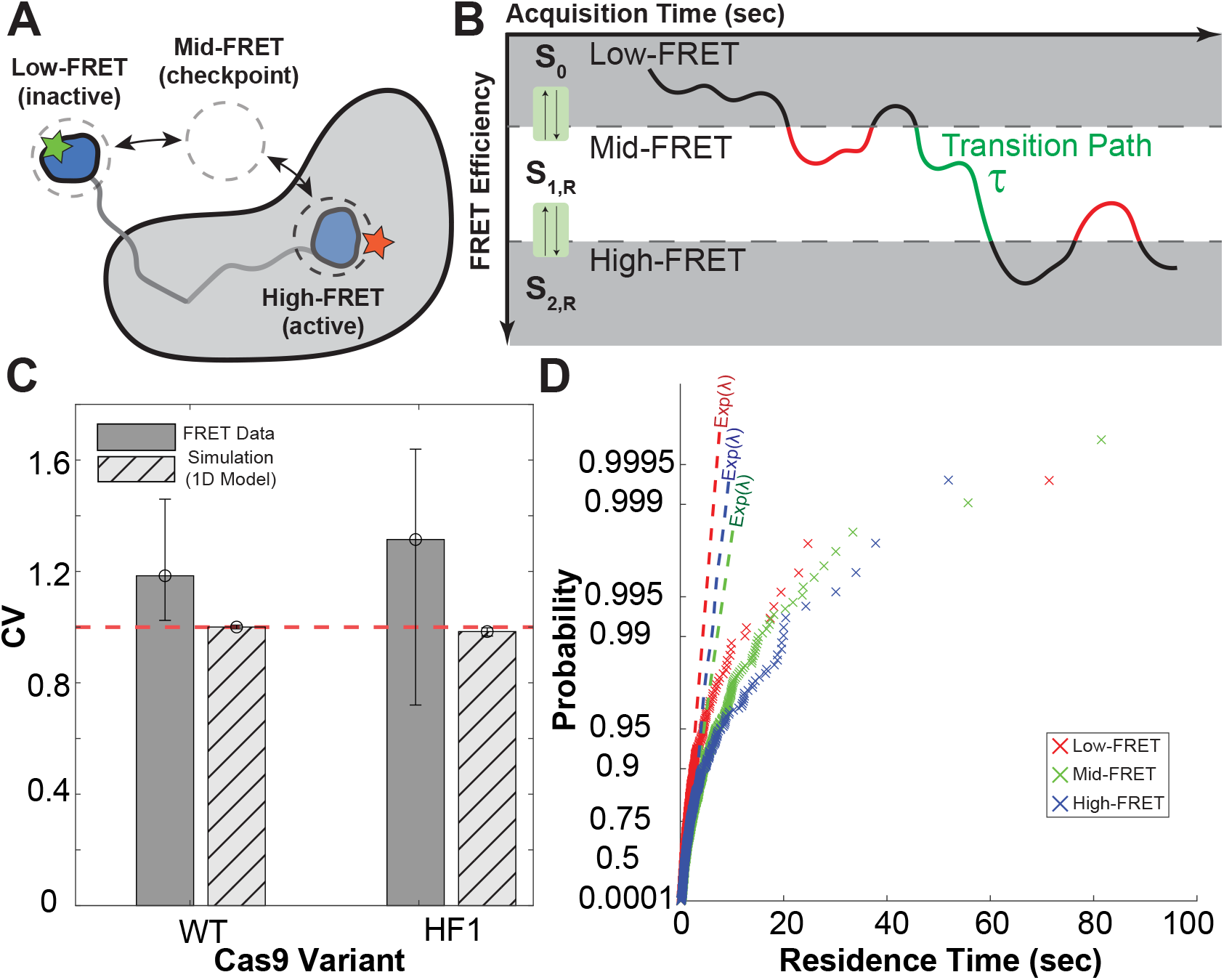
The Cas9 HNH domain was identified as a potential contributor to cleavage specificity and found in smFRET studies to have three possible conformational states corresponding to low-, mid-, and high-FRET configurations (A). The HNH domain dynamics can yield insights into the structure of the Cas9 kinetic network. The statistics of HNH domain transition paths (B) which are the time required for the HNH domain to transit from the low-FRET state to the high-FRET state (green path) can help differentiate between possible model structures. The smFRET dataset suggests that the distribution of transition paths is broad, with CVs greater than unity for both WT and HF1-Cas9 variants (C). The broad distribution of transition paths suggests that the underlying kinetic network must be multi-dimensional, with a 1D kinetic model (boxed network in Fig. 1A) being incapable of producing transition path distributions with CVs greater than one (C, hatched bars). Additionally, the distribution of residence times for the three HNH domain states show significant deviations (*p* ≪ 0.05, Kolmogorov-Smirnov test) from reference exponential distributions (dashed reference lines, passing through the 1st and 3rd quartiles of the sample data), which we expect only for multi-dimensional models (D).

Recent theoretical results have shown that the statistics of single-molecule transitions can reveal deep insights into the underlying thermodynamic landscape. In particular, the statistics of transition path times can help differentiate between different kinetic model structures [24]. Here, a transition path occurs when Cas9 in the low-FRET HNH domain successfully transits through the mid-FRET state to the high-FRET state (Fig. 2B).

We identified all such transition paths in the smFRET dataset for both WT-Cas9 and HF1-Cas9. For each variant, the distribution of transition path times was broad, with coefficients of variation (CVs) greater than one (Fig. 2C). The CV of a sample is defined as the standard deviation (*σ*) scaled by the mean (*µ*), CV = *σ*/*µ*. When the CV is greater than one, it indicates that the underlying thermodynamic landscape must be multi-dimensional [24]; in essence, multiple paths must exist for the system to transit from point A to point B. In the context of Cas9, multi-dimensionality implies that the simple 1D model is insufficient to properly model Cas9 behavior. As expected, Monte Carlo simulations of the 1D model parameterized with rate constants from Liu et al. [18] produce transition path CVs less than or equal to one most of the time for both WT-Cas9 and HF1-Cas9 (Fig. 2C).

We can also examine the residence times distribution for each HNH domain state to gain insight into the proper model structure. The built-in MATLAB^®^ `probplot` function generates probability plots [32] that allow sample data to be compared against the quantiles (converted to cumulative probabilities) of a reference probability distribution. If Cas9 binding and cleavage were a 1D process, as in the boxed network in Figure 1A, the distribution of residence times for the low and high-FRET states would be exponential. Thus, the sample data would be roughly linear when plotted against quantiles from exponential distributions with identical means (2D). However, the experimental data shows substantial deviations from the exponential distribution for all three HNH domain states, which further suggests that the underlying kinetic network must be multi-dimensional.

### C. A 2D kinetic model can match experimental Cas9 datasets

The simplest multi-dimensional kinetic network that can describe Cas9 behaviors is a 2D network in which target recognition (PAM-binding, R-loop formation/collapse) can occur before or after the HNH domain transitions (Fig. 3). We implement this model similarly to the 1D model, with Cas9 selecting between an on-target and off-target substrate. Unbound Cas9 can exist in either the low-FRET or mid-FRET state, as HNH domain transitions were detected when Cas9 is only bound to the guide-RNA [1, 3].

**Figure 3.**
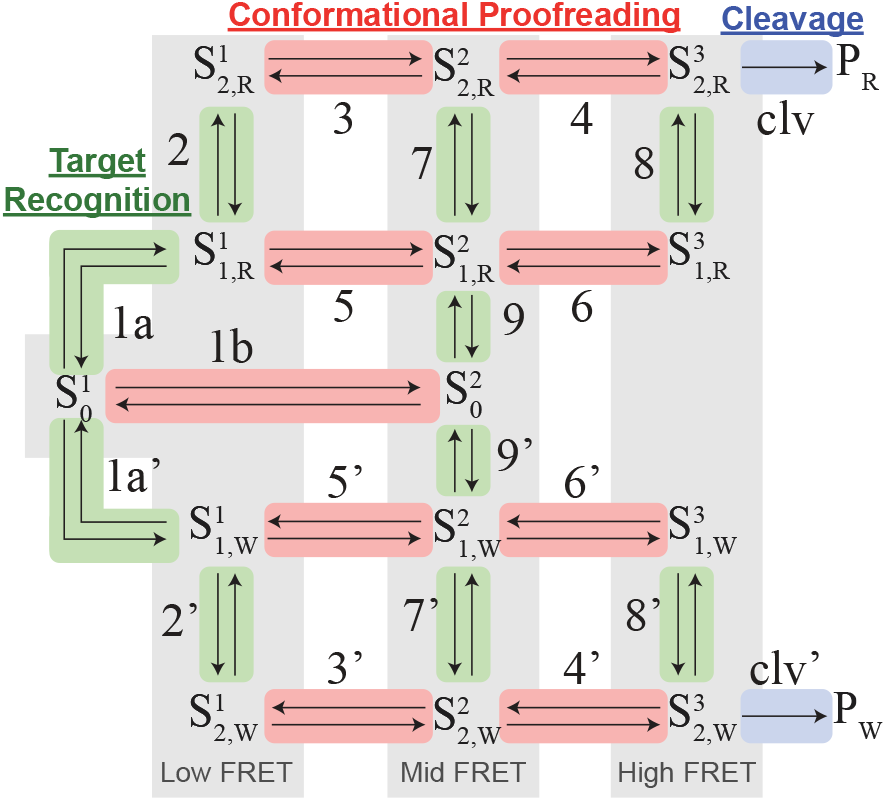
A 2D model of Cas9 target binding and cleavage can be constructed by allowing the R-loop formation/collapse dynamics to occur orthogonally to the movements of the HNH domain. Reactions can be grouped into three distinct sets of processes, including those involved in target recognition (green), conformational proofreading (red), and substrate cleavage (blue). The individual model states are binned into low-FRET 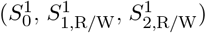 mid-FRET 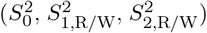 and high-FRET 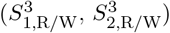 states.

We fitted the 2D model to various datasets, including the HNH domain smFRET dataset and a high-throughput dataset of Cas9 substrate cleavage rates and binding affinities. The fitted model is high quality, with almost all on-target data points fitted within ±25% margin (Fig. 4A).

**Figure 4.**
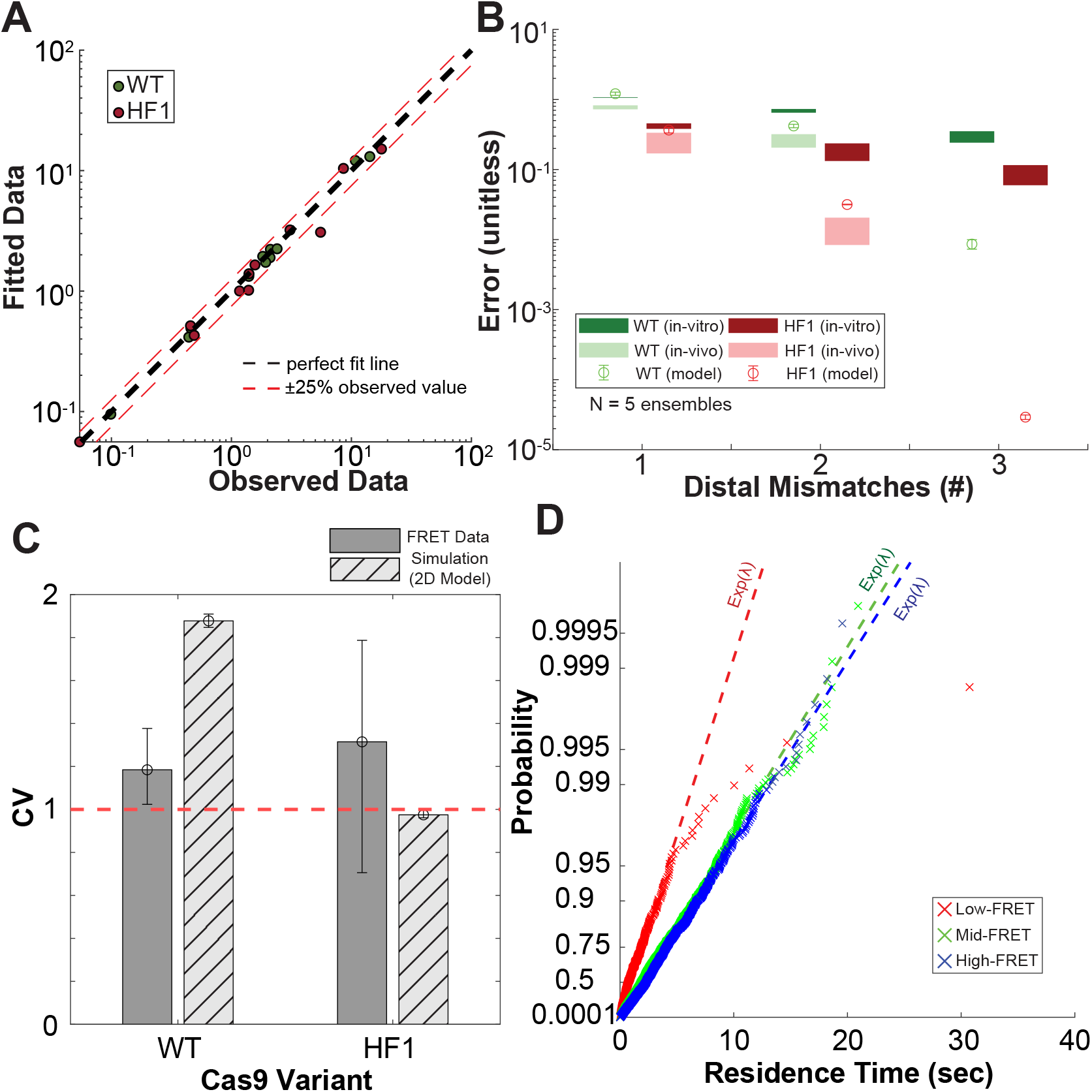
The 2D model can be well fitted to a dataset consisting of high-throughput binding and cleavage measurements as well as HNH domain transition statistics derived from smFRET (A). The fitted model produces off-target cleavage error predictions that are qualitatively consistent with experimental measurements in both in-vitro and in-vivo settings (B). Monte Carlo simulations of the fitted model produce transition path distributions consistent with the experimental dataset (C). Further, Monte Carlo simulations with the fitted 2D model produce non-exponential HNH domain residence time distributions that deviate from the reference exponential distribution (dashed reference lines, passing through the 1st and 3rd quartiles of the sample data), which are consistent with the experimental smFRET dataset (D).

The model produces cleavage error predictions that are qualitatively consistent with both *in-vivo* and *in-vitro* datasets (Fig. 4B). Quantitative differences between the experimental datasets and the model predictions are likely due to limitations in the available fitting data. Specifically, the model is fitted to *in-vitro* data. Thus, the data does not account for cellular factors that could affect Cas9 specificity, such as the presence of histones or active DNA remodeling. Furthermore, due to the low throughput of the FRET experiments, we were required to assume that the dynamics of the HNH domain would depend solely on the number of PAM-distal mismatches; thus, the *N* mismatch substrates in the data set were fitted to have the same HNH domain dynamics as the corresponding substrate in the smFRET data set.

We additionally performed Monte Carlo simulations with the fitted model to check for agreement with the smFRET dataset; the Monte Carlo simulations were performed for the single-substrate case (no off-target substrate present, 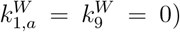. The simulations produce transition path CVs that are greater than one for WT-Cas9, which is consistent with the experimental data (Fig. 4C). The HF1-Cas9 transition path CV is less than one, but still remains within the 95% confidence interval obtained from the experimental dataset.

Monte Carlo simulations with the fitted model produce transition path CVs that are greater than one, which is consistent with experimental data. The simulations of the WT-Cas9 model also produce residence-time distributions that are non-exponential (for the low-FRET state, *p* ≈ 10^−4^, Kolmogorov-Smirnov test), which is consistent with what is observed in the experimental data (Figs. 2D and 4D).

### D. HNH domain dynamics are a key contributor to specificity

Next, we attempted to use our model to understand the relative importance of key Cas9 processes for substrate selection specificity. Previous work has suggested that processes such as the dynamics of the HNH domain [1, 3], R-loop formation and collapse dynamics [8, 27], and the rate of substrate cleavage [18] all contribute to the accuracy of substrate selection. Each of these processes is represented as distinct transitions in our model, which allows us to gauge their relative importance quantitatively.

We performed a global parameter swapping routine to determine which processes in the HF1 high-fidelity Cas9 model led to the greatest improvement in the substrate selection error for wild-type Cas9. We focused on the HNH domain dynamics (reactions 3, 4, 5, 6 in Fig. 3A) and the R-loop formation and collapse dynamics (reactions 2, 7, and 8 in 3). The microscopic substrate cleavage rate was assumed to be identical across both Cas9 variants and substrates and, therefore, is not relevant to this analysis.

The native WT-Cas9 and HF1-Cas9 (ensemble average) cleavage error predictions for the distal-mismatch substrates are shown in blue and purple in Fig. 5; the trend is qualitatively consistent with experimental measurements of Cas9 off-target accuracy [15]. We see that while both the HNH domain dynamics (yellow) and R-loop dynamics (orange) lead to a reduction in WT-Cas9 cleavage error, the HNH domain dynamics have a larger effect (Fig. 5A). From this, we conclude that the HNH domain dynamics are the primary contributor to specificity for distally mismatched substrates.

**Figure 5.**
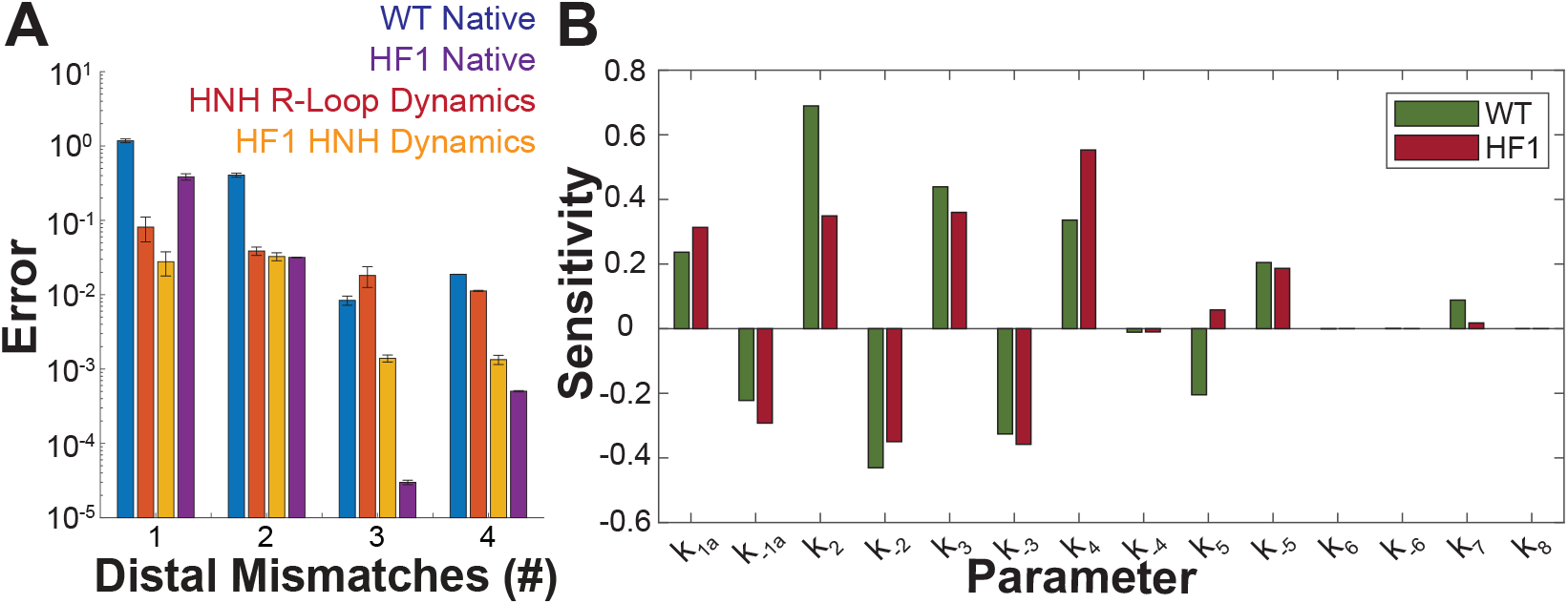
Globally swapping sets of parameters corresponding to R-loop formation/collapse as well as HNH domain transitions between the WT and HF1-Cas9 models can yield insights into the relative importance of each process on specificity. The swapped models suggest that the HF1-Cas9 HNH domain dynamics (A, yellow bars) have a greater impact on specificity than HF1-Cas9 R-loop formation/collapse dynamics (A, orange bars). Additionally, the low-FRET R-loop formation/collapse rates and the HNH domain transition rates in the R-loop state are predicted to have the greatest effect on on-target cleavage speed (B).

We additionally performed a local sensitivity analysis to identify which model parameters are predicted to have the greatest effect on the on-target cleavage speed, *V* = 1/*τ*. For both the WT and HF1 variants, the R-loop formation and collapse rates (*k*_2,−2_) and HNH domain transitions in the R-loop state (*k*_3,−3_, *k*_4,−4_) have the most significant effect on the speed of on-target cleavage (Fig. 5B).

### E. There are no intrinsic speed-specificity trade-offs for Cas9

Finally, we used our model to examine speed-specificity trade-offs in Cas9. Previous experimental studies identified a clear trade-off in on-target cleavage efficiency (measured as on-target indel formation rate) and off-target cleavage specificity (measured as the off-target indel formation rate relative to the on-target) [15]. Our model can be used to compute analogous quantities to gauge the effects of distinct thermodynamic parameter perturbations on Cas9 specificity and efficiency. The model can compute the off-target cleavage specificity directly, however we must use the speed of on-target cleavage as a proxy for on-target cleavage efficiency.

While the model predicts the existence of a speed-specificity trade-off when the barrier for substrate cleavage is perturbed (Fig. 6A), there is no such trade-off observed for perturbations in the R-loop formation barrier (Fig. 6B) or for perturbations in the HNH domain transition barriers in the fully formed R-loop state (Fig. 6C,D). Overall, the lack of a clear speed-specificity trade-off for the HNH domain dynamics or R-loop formation and collapse dynamics suggests no biophysical limitation to developing high-specificity, high-efficiency Cas9 variants. ‘

**Figure 6.**
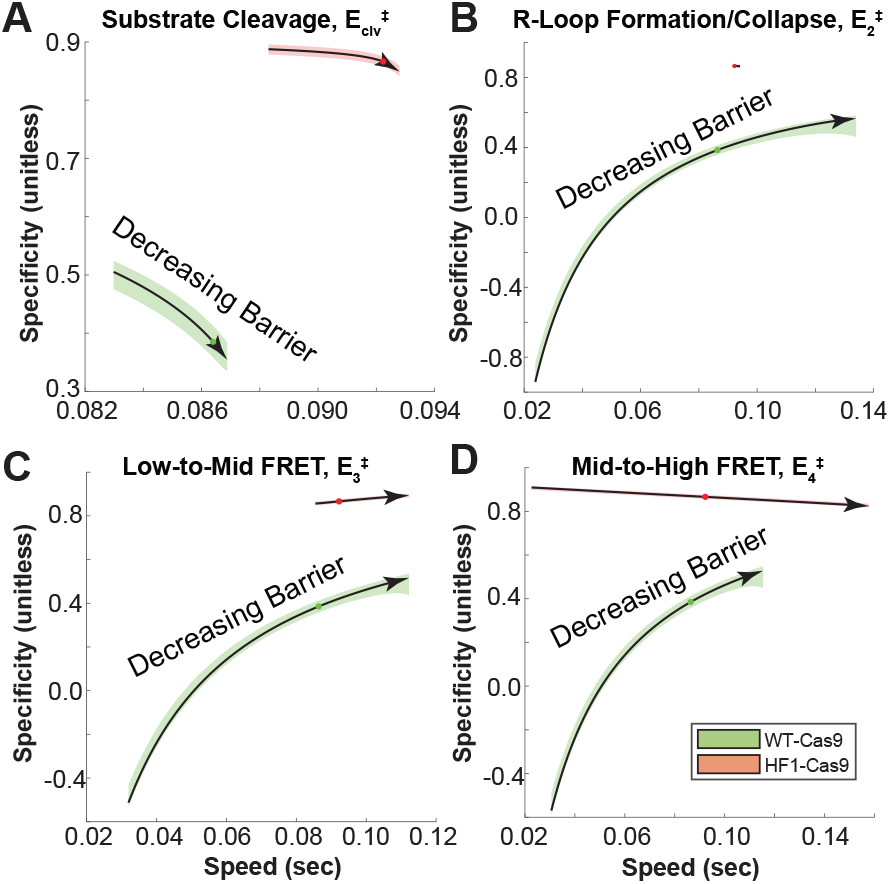
Perturbing free-energy barriers in the fitted model allows us to identify potential parameter regimes that would allow Cas9 to achieve both high on-target cleavage efficiency and low off-target cleavage (high-specificity). While there is a clear trade-off for perturbations in the substrate cleavage barrier (A), for R-loop formation/collapse (B) as well as HNH domain transitions in the fully formed R-loop state (C, D) there is no appreciable trade-off, with Cas9 being able to increase on-target cleavage speed with minimal impacts on cleavage specificity.

## III. DISCUSSION

Here, we have presented a model of CRISPR-Cas9 substrate selection and cleavage. The model fits diverse experimental data sources, including high-throughput measurements of Cas9-mediated DNA binding and cleavage [14], as well as single-molecule measurements of the Cas9 HNH domain dynamics when bound to PAM-distally mismatched DNA substrates [1, 3]. The fitted model reproduces trends observed in the literature, such as the effective mid to high-FRET transition rate, which was noted to be higher in WT-Cas9 than the HF1-Cas9 variant [1].

The HF1-Cas9 [17] high-fidelity Cas9 variant was developed under the “excess energy” hypothesis, which requires that the Cas9-DNA binding affinity be decreased to improve the accuracy of substrate selection. However, we have shown that changes in binding affinities are insufficient to affect the accuracy of Cas9 substrate selection, which is consistent with both experimental measurements [1] and prior theoretical results for generic reaction networks [20]. Instead, any change in the accuracy of Cas9 requires changes in the transition state energies (i.e. kinetics) of substrate binding and cleavage. Our model predicts that the HF1-Cas9 variant is more likely to get stuck in the mid-FRET, checkpoint state compared to the wild-type variant, as the dissociation constant for entering the checkpoint state with a fully-formed R-loop (*K*_*D*_ = *k*_−3_/*k*_3_) is much higher than for wild-type Cas9. Additionally, when HF1-Cas9 is targeted to off-target substrates with a single distal mismatch, the enzyme tends to get stuck in the low-FRET, full R-loop state (*S*_2,1_) compared to WT-Cas9; the fraction of Cas9 proteins in either of the two mid-FRET states are predicted to be roughly equal between WT-Cas9 and HF1-Cas9.

A previous model developed by Klein and Eslami-Mossallam [4, 16] considers Cas9 substrate binding and cleavage as a 1D birth-death process in which the R-loop grows a single base at a time until fully formed. Under this model, any HNH domain movements would be required to occur either simultaneously with R-loop formation or as an additional step following the formation of the full R-loop. In contrast, statistical analysis of the Cas9 HNH domain dynamics (from smFRET) has revealed a broad distribution of HNH domain transition path times, which suggests that the HNH domain dynamics are best modeled as processes independent of R-loop formation and collapse. Overall, this implies that the HNH domain is not tightly mechanically coupled to the formation of the R-loop, and should instead be modeled as separate, distinct reaction steps that can occur in the presence of an incomplete or partially formed R-loop.

Structural studies by Pacesa et al. show that the HNH domain requires partial R-loop formation (at least through the “seed” region) to undock [22]. In our model, because we allow for differences in PAM unbinding rates between the on and off-target substrates, our PAM-bound state is more akin to an early (or “seed” region) R-loop state. In this case, we expect the HNH domain to still be capable of movement in either of the bound states. Regardless, the observation of transitions between the low-FRET and mid-FRET states in the absence of DNA substrate suggests that partial R-loop formation is not a strict requirement to observe HNH domain movements [3].

Additionally, separate smFRET studies by Singh et al. found that R-loop formation occurs as a two-step process with an initial “sampling” state followed by the formation of a fully constituted R-loop [26]. The observation that R-loop formation occurs as a two-step process is consistent with our model as we explicitly require binding to occur in two steps. Our fitted model also suggests that HF1-Cas9 is more likely to reject off-target substrates from the initially bound, partial R-loop state than wild-type Cas9. This appears to be consistent with later findings by Singh et al., which showed that the R-loop formation and collapse dynamics of HF1-Cas9 are more sensitive to the presence of PAM-distal mismatches [27].

Our model produces cleavage error predictions for PAM-distally mismatched substrates that are qualitatively consistent with both *in-vivo* and *in-vitro* measurements of Cas9 cleavage error. However, some quantitative differences remain between model predictions and experimental measurements. Namely, our model predicts cleavage errors that are intermediate between the *in-vivo* and *in-vitro* measurements for a given number of distal mismatches. There are multiple possible causes for this. First, due to limited smFRET data (which is only available for five substrates, and is missing for the wild-type 1bp and 2bp mismatch substrates), we were forced to assume that only the number of PAM-distal mismatches would affect the HNH domain dynamics. Therefore, for each substrate in the high-throughput dataset, we assumed that the HNH domain dynamics to be identical to the corresponding smFRET substrate based on the number of distal mismatches. In reality, the precise identity and ordering of PAM-distal mismatches, as well as the presence of PAM-proximal mismatches, would likely affect the behavior of the HNH domain. Further, even with the wild-type 1bp and 2bp substrates displaying steady-state FRET histograms that are largely similar to the on-target substrate [1], it is possible that the precise HNH domain dynamics vary for these conditions and could affect the fitted transition rates. Second, the fitting data we use to parameterize the model comes exclusively from *in-vitro* experiments. In the *in-vitro* context, multiple factors that could affect Cas9 cleavage are absent, such as the presence of histones in nucleosomes [10] and DNA stretching due to active processes such as RNA transcription and nucleosome remodeling [21]. The influence of these factors could affect the accuracy of Cas9 and should be accounted for in future models, especially as they pertain to genome therapies.

Importantly, our model reveals the relative importance of different Cas9 biophysical processes in substrate selection. Prior work has focused on the HNH domain dynamics [1, 3], R-loop formation and collapse dynamics [8, 27], and substrate cleavage rate [18] as possible contributors to substrate specificity. Our model suggests that while both the HNH domain dynamics and R-loop formation/collapse dynamics are important for substrate selection, the dynamics of the HNH domain are the most significant factor controlling off-target cleavage for PAM-distally mismatched substrates.

Our work shed important insights into trade-offs in Cas9 engineering. Previous work by Kim et al. has identified trade-offs between Cas9 substrate selection fidelity and on-target cleavage efficiency [15], in which the efficiency of on-target cleavage tends to decrease as the ability to reject off-target substrates increases. Examining different perturbations in our model suggests that it is generally possible for Cas9 to avoid this trade-off. While our model cannot predict on-target cleavage efficiency as defined by Kim et al., instead predicting the speed of on-target cleavage, it suggests that further engineering should allow for high-efficiency, high-specificity Cas9 variants. Indeed, recent work using generative artificial intelligence (GenAI) models has lead to the development of an artificial Cas9-like enzyme (OpenCRISPR-1) that appears to achieve greater off-target specificity than wild-type Cas9 while maintaining wild-type-like on-target cleavage efficiency [23]. The development of the OpenCRISPR-1 variant is consistent with our finding that there is no intrinsic barrier preventing high-efficiency, high-specificity Cas9 variants. However, it will still be important to test the OpenCRISPR-1 variant on a much larger set of off-target substrates to get a more accurate sense of the specificity enhancement relative to wild-type Cas9.

## IV. METHODS

### A. Single-Molecule FRET Data and Processing

Single-molecule FRET traces for wild-type, HF1, and eSpy(1.1)-Cas9 were obtained from the study of Chen et al. 2017 [1, 3]. Chen et al. acquired smFRET data for each Cas9 variant bound to either a fully matched, on-target substrate or PAM-distally mismatched substrates with one to four mismatches. Each trace in the dataset consists of the Cy3 (donor) and Cy5 (acceptor) signal intensity (in a.u.) over time for an individual Cas9-DNA particle with FRET labels attached to the Cas9 HNH and REC1 domains [1]. The Chen dataset was acquired using a camera with a 10Hz acquisition frequency [1]; therefore, the time resolution of the data is limited to 100 milliseconds. We were unable to obtain FRET traces corresponding to the WT-Cas9 1bp and 2bp mismatch data; we make the assumption that the WT HNH domain dynamics for the 1bp and 2bp mismatch cases will be similar to the fully-matched, on-target case because they have similar steady-state FRET histograms [1]. Therefore, we re-use the on-target FRET data for the WT-Cas9 1bp and 2bp conditions.

The FRET traces were preprocessed prior to statistical analysis. First, the donor and acceptor signal intensity traces were smoothed using a 3rd-order Savitsky-Golay filter with a window size of 11 timesteps. The MATLAB^®^ function `sgolayfilt` in the Signal Processing Toolbox was used to implement the filter. The timepoint corresponding to acceptor photobleaching was identified using the `photobleach index` function in the ebFRET package [30, 31]; individual traces were then truncated at the point of photobleaching. The dataset was then filtered to select traces displaying anti-correlated donor and acceptor signals (Pearson’s *ρ* < −0.5), which is indicative of a true FRET signal. Traces shorter than 50 frames (5 seconds) after truncation were discarded. For each trace, the effective FRET signal was computed as the ratio

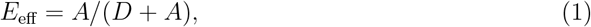

where *D* and *A* are the donor and acceptor intensities, respectively.

### B. FRET State Discretization

The processed smFRET traces were discretized using the empirical Bayes hidden Markov model (HMM) tool ebFRET [30, 31]. The ebFRET tool implements a linear HMM that can be fitted simultaneously to large datasets by learning a distribution of model parameters to account for variability between molecules due to slight differences in experimental conditions, photophysical effects, and other factors [30]. Discretizing using an HMM helps to assign states to each point in the smFRET trace while minimizing the effects of signal noise. We fitted the traces in ebFRET to a six-state HMM with default settings. Prior to fitting, outlier FRET intensities within each trace were clipped to the range [−0.2, 1.2]; traces with more than 10 outliers were discarded. The discretized states were then assigned to the low-FRET (inactive), mid-FRET (checkpoint), and high-FRET (active) HNH domain states by a simple thresholding procedure with thresholds set based on the mean FRET efficiencies reported by Chen et al. [1]. The threshold *T*_*ij*_ between adjacent states *i* and *j* was set as the midpoint between the FRET efficiencies:

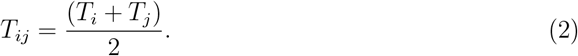

### C. HNH Domain Residence Time Statistics

HNH domain state residence times were computed from the FRET traces using custom scripts in MATLAB^®^. After each detected transition, the time until the next transition was computed and taken to be the dwell time *τ*.

HNH domain state residence times were computed from each discretized trace in the smFRET dataset. The mean residence time for each FRET state *i* is an average

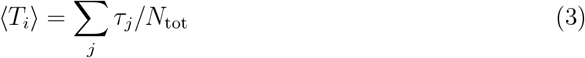

where each *τ*_*j*_ represents the time the system stays in state *i* prior to a detected state transition (residence time), and *N*_*tot*_ is the total number of detected events. For each trace, we discard the first and last detected transitions as we do not know the proper transition start and end times for these transitions.

### D. HNH Domain Steady-State Occupancy

The probability of observing each HNH domain state at steady-state was calculated from the smFRET dataset as an average over the final timepoint for each trace. We assume that each trace is sufficiently long for the ensemble of traces to achieve a steady state. The steady-state probabilities 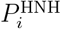 are computed as

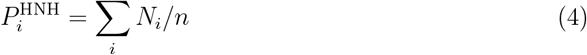

where *n* is the total number of traces and *N*_*i*_ is the number of traces that end in state *i*.

### E. HNH Domain Transition Paths

Transition paths are computed from smFRET traces by tabulating all transitions from the mid-FRET (checkpoint state) into the high-FRET (active state). The initial transition path for a given FRET trace is discarded if the trace starts in the mid-FRET state, as there is no way to determine how long the molecule has resided in the mid-FRET state. After a transition occurs, the FRET trace must return to the low-FRET state before additional transition paths are counted.

### F. High-Throughput Data Preprocessing

High-throughput measurements of WT-Cas and HF1-Cas9 substrate binding and cleavage were obtained from Jones et al. [14]. For each substrate in the dataset, we computed the effective cleavage mean first-passage time from the measured cleavage rate *k*_*i*_ as

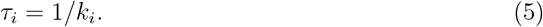

The effective cleavage error for each substrate can then be computed as a ratio of mean first-passage times

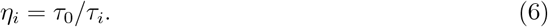

where *τ*_0_ is the cleavage mean-first passage time for the on-target substrate.

The dataset was filtered to select substrates that possess a valid NGG PAM sequence, as well as substrates for which the effective cleavage error *η*_*i*_ is less than one. The high-throughput dataset was preprocessed using a custom script in Python 3.

### G. Transition State Theory

The transition rate constant for each transition *i* → *j* can be computed via transition state theory as

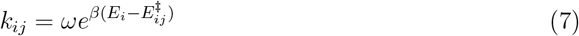

where *E*_*i*_ is the free-energy of state *i* and 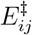 is the free-energy barrier for the *i* → *j* transition. The parameter *ω* is a pre-exponential frequency factor and *β* = 1/(*K*_*B*_*T*), where *K*_*B*_ is the Boltzmann constant.

### H. Forward Master Equation

The forward chemical master equation is defined by the system of differential equations

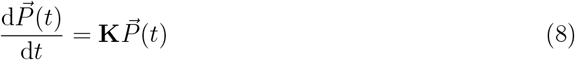

where 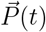 is a vector of state probabilities and the matrix **K** encodes the state transitions. The system of equations is subject to the normalization constraint

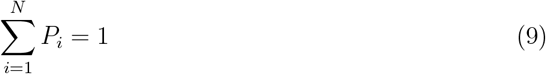

which ensures that each *P*_*i*_ can be interpreted as a probability. We can obtain the steady-state probability for each system state with the condition

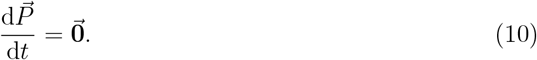

### I. Backward Master Equation

The backward chemical master equation is defined as a system of differential equations

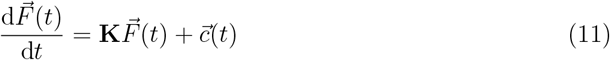

where each *F*_*i*_(*t*) is a probability distribution giving the probability of reaching some terminal state (e.g.: the cleaved substrate) at time *t*, conditioned on the system starting in state *i* at time *t* = 0. The matrix **K** is a transition matrix that encodes the state transitions of the model, and 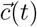 is a vector of source terms that encodes the initial conditions (e.g., *F*_*q*_(0) = 1 if state *q* is a terminal state). Note that the transition matrix **K** is distinct from the transition matrix used in the forward master equation.

The system of differential equations defining the backward master equation can be transformed into Laplace space to yield

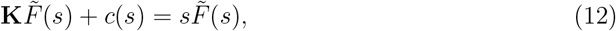

which can be solved algebraically for each 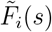.

Critical properties of the system can then be obtained from 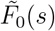, which represents the system starting in the unbound, low-FRET state. The system splitting probabilities *π*_*R*_ and *π*_*W*_ can be obtained by taking the limit of 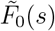 as the Laplace variable s goes to zero:

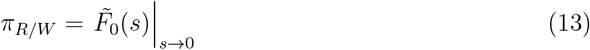

The effective substrate selection error (cleavage error) can then be defined as a ratio of splitting probabilities:

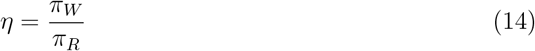

The backward master equation can additionally used to obtain information on the system dynamics, such as the mean first-passage time (MFPT, *τ*) to on-target (R) substrate cleavage. The MFPT is obtained by computing

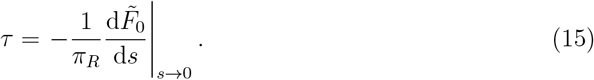

We can also compute the effective cleavage speed as the inverse of the *τ*,

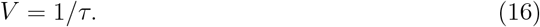

### J. Dissociation Constant and Binding Affinities

The apparent Cas9-substrate dissociation constant (*K*_*D*_) can be computed from our kinetic modeling framework by computing the steady-state bound fraction, *P*_bound_ for varying substrate concentrations (*x*) using the forward master equation [14]. The bound fraction is defined as the sum of all bound states or equivalently

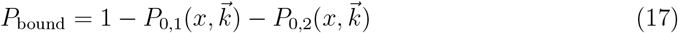

where *P*_0,1_ and *P*_0,2_ are the low-FRET and mid-FRET unbound states.

The dissociation constant *K*_*D*_ is equivalent to the concentration of substrate at which the available Cas9-gRNA is half-saturated, i.e. *P*_*bound*_ = 1/2. We can obtain 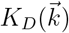 as a function of model parameters by solving the equation

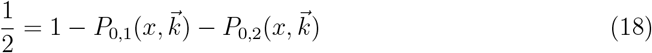

for the concentration *x* = *K*_*D*_, where 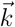 is a vector of model parameters.

The normalized binding affinity (Δ*ABA*) for off-target substrates is computed numerically as the ratio of the effective dissociation constants *K*_*D,R*_ and *K*_*D,W*_, i.e. Δ*ABA* = *K*_*D,W*_ /*K*_*D,R*_.

### K. HNH Domain Residence Time Statistics

We use the framework of Gopich and Szabo [9] to compute residence time distributions for each of the HNH domain FRET states. For each HNH domain FRET state, we consider only the DNA bound states (i.e. ignoring states 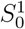 and 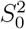).

The matrix **V** encodes the transitions that enter the meta-state of interest (e.g. ‘low-FRET state’). The entry probability is computed from the steady-state solution of the system as

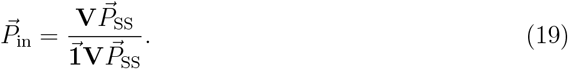

The matrix **U** encodes the transitions that exit the meta-state of interest. We can solve for the generating function for the distribution of exit times by solving

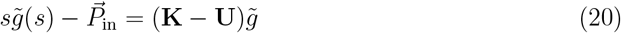

where 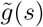 is the generating function in Laplace space.

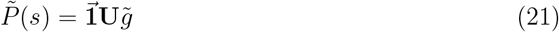

The raw moments of the residence time distribution can be computed from 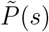 bydifferentiating

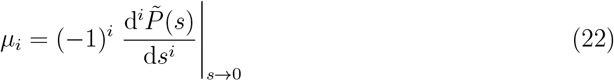

and taking the Laplace variable *s* → 0.

The residence time mean is identical to the first raw moment:

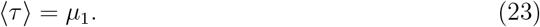

The residence time variance is obtained as

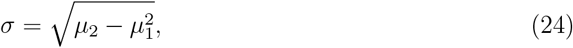

from which we can then obtain the residence time CV:

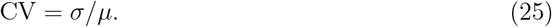

### L. Model Optimization

We parameterized the model using a two-stage fitting procedure. First, we fitted the model to the WT-Cas9 and HF1-Cas9 on-target data along with data from a 4bp mismatched substrate. After the initial fitting, we then fixed the on-target parameters and fitted the model to the remaining off-target substrates in the dataset individually. For both fitting stages, we used the built-in particle swarm optimizer (`particleswarm`) available in MATLAB^®^ Global Optimization Toolbox. All model fitting runs were performed on the Rice University NOTS high-performance computing cluster in MATLAB^®^ R2021a.

Additional details on the model implementation and fitting procedure are given in the Supplemental Information.

### M. Sensitivity Analysis

Local sensitivity analysis is performed by computing logarithmic gains as

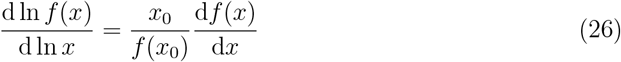

where *x* is a model parameter of interest and *f*(*x*) is some model output. The terms x_0_ and *f*(*x*_0_ are the initial parameter value and the model response at that point.

### N. Monte Carlo Simulation

Stochastic simulations of the model were performed using a custom implementation of the Gillespie algorithm [7] in MATLAB^®^ R2021a. Simulations were performed for *N* = 1000 trajectories, with each trajectory simulated for *t* = 100 seconds. At the beginning of each trajectory we simulated an additional 800 seconds of burn-in time to ensure that the simulation results were unbiased by the initial conditions.

### O. Cleavage Specificity

The average cleavage specificity across all off-target substrates in the dataset for each variant is computed from the average cleavage error

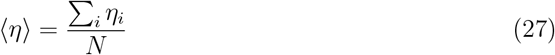

as

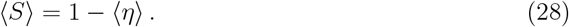

### P. Probability Plots

Probability plots comparing a data sample against a theoretical reference distribution are generated using the built-in MATLAB^®^ function `probplot`. The `probplot` function generates a probability plot in which the quantiles from the theoretical distribution are rescaled and converted to the equivalent cumulative probability. The reference line generated by `probplot` is generated from the theoretical distribution and is set to pass through the 1st (25%) and 3rd (75%) quartiles of the sample data.

### Q. Statistical Tests

Kolmogorov-Smirnov tests were performed using the built-in MATLAB^®^ function `kstest` with the significance level α = 0.01. Sample distributions were compared against exponential distributions set to have the same mean as the sample.

## VI. AUTHOR CONTRIBUTIONS STATEMENT

A.D.H. and O.A.I. conceived the study, A.D.H. conducted the experiment(s), A.D.H. and O.A.I. analysed the results. A.D.H. and O.A.I. wrote and reviewed the manuscript.

## VII. ACKNOWLEDGEMENTS

We wish to thank Yavuz Dagdas and Jennifer Doudna for providing us with their smFRET dataset. Additionally, we wish to thank A. Kolomeisky, G. Bao, P. Murphy, Z. Diao, and R. Butcher for useful discussions. We also wish to thank G. Bao and D. Makarov for useful comments on the manuscript. The research was supported by the Welch Foundation (Grant C-1995 to OAI).

This work was supported in part by the Big-Data Private-Cloud Research Cyberinfrastructure MRI-award funded by NSF under grant CNS-1338099 and by Rice University’s Center for Research Computing (CRC).

## Supplemental Information

### 1. Model Implementation

#### 1.1 Model Structure

We model CRISPR-Cas9 substrate selection as a two-branch process in which unbound Cas9 (assumed to be preloaded with a guide RNA) can select between two possible substrates: a fully matched, on-target substrate (R) and a mismatched, off-target substrate (W). The model structure is agnostic to the number and relative positions of mismatches between the Cas9 guide RNA and the target site DNA.

The transitions between model states *S*_*i*_ and *S*_*j*_,

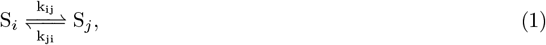

have rate constants defined in terms of transition state theory. Each rate constant is defined in terms of the energy of the starting state (*E*_*i*_) and the corresponding reaction barrier 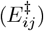:

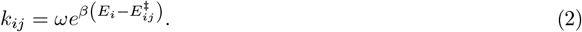

The term *ω* is a prefactor that ensures that the rate constant has the proper units (*s*^−1^), while *β* = 1*/*(*K*_*B*_*T*) sets the energy scale.

The on-target and off-target model branches are related through discrimination factors *f*_*i*_ = *k*_*i,W*_ */k*_*i,R*_, which fix the ratio between the on-target and off-target rate constants for reaction *i*. When the discrimination factor *f*_*i*_ = 1, it implies that there is no discrimination between on- and off-target substrates for reaction *i*.

#### 1.2 Model Fitting

The model objective function for fitting is defined as

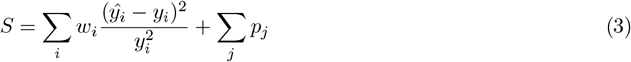

where 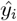 and *y*_*i*_ are the predicted and observed data points, respectively. The constant *w*_*i*_ is a weighting factor that allows us to scale the relative contribution of each data point to the overall fit. Additionally, we include a set of penalty terms *p*_*i*_ that enforce various constraints on the model parameters (such as upper bounds).

The model is fitted to the experimental data using particle swarm optimization (PSO) using the MATLAB^®^ Global Optimization toolbox function `particleswarm`. The initial conditions for the particle swarm were set based on a previous best fit model in which the model rate constants were obtained by a direct fit, instead of fitting the corresponding state energies (*E*_*i*_) and reaction barriers 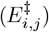.

#### 1.3 Model Assumptions and Constraints

The model is fitted using several assumptions. First, we assume that the R-loop formation and collapse rates (*k*_2_, *k*_−2_, *k*_7_, and *k*_8_) have the same discrimination factors for both the WT and HF1-Cas9 variants. In essence, we assume that the protein modifications made by Kleinstiver et al. [3] do not materially affect the rates of R-loop formation and collapse. Second, we similarly assume that the PAM unbinding rate *k*_−1,*a*_ is unchanged across Cas9 variants. Third, we assume that the initial PAM binding rates (*k*_1,*a*_ and *k*_9_) are not only identical between the on- and off-target substrates but that the on-target rates are identical across Cas9 variants. We make a similar assumption for the microscopic cleavage rate, *k*_*clv*_.

We enforce “soft” upper bounds for each rate constant to ensure that they take on physically realistic values after fitting. For each parameter *k*_*i*_, we apply a penalty of the form

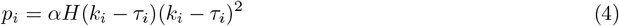

where *H*(*x*) is the Heaviside step-function and *τ*_*i*_ is the upper bound for the given parameter. The constant *α* is a scaling factor that allows us to vary the magnitude of the penalty; we generally set *α* = 1000 to ensure that the upper bounds are always satisfied.

Finally, we fix the Cas9 HNH domain transitions in the unbound state to the values reported by Dagdas et al. (*k*_1,*b*_ ≈ 0.29s^−1^, *k*_−1,*b*_ ≈ 4.2s^−1^, [1]). Since these reactions occur in the unbound state, there are no corresponding discrimination factors to consider. We enforce these rates by again applying a soft penalty term of the form

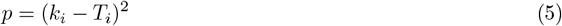

where *T*_*i*_ is the target value for the rate constant *k*_*i*_.

#### 1.4 Fitting Substrate Cleavage Rates

We use NucleaSeq data published by Jones et al. [2] to obtain the effective substrate cleavage rates for different Cas9 variant-substrate pairings. The effective cleavage rates are then used to estimate the mean first-passage time (MFPT) for substrate cleavage as

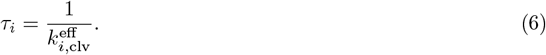

In the Jones experimental setup, the effective limit of detection is approximately 6e4 seconds and is limited by the length of the experimental time-course. Any Cas9-substrate pairs that cleave on timescales longer than the experimental timescale cannot be effectively estimated, as there will be very few cleavage events to detect. To account for these substrates, when fitting the model to the cleavage MFPT data we apply a penalty only if the model predicted MFPT is slower than the detection limit, i.e.

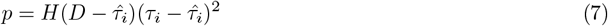

where *D* is the limit of detection and *H*(*x*) is the Heaviside step function.

